# Simultaneous isotope dilution quantification and metabolic tracing of deoxyribonucleotides by liquid chromatography high resolution mass spectrometry

**DOI:** 10.1101/454322

**Authors:** Rostislav Kuskovsky, Raquel Buj, Peining Xu, Samuel Hofbauer, Mary T Doan, Helen Jiang, Anna Bostwick, Clementina Mesaros, Katherine M. Aird, Nathaniel W. Snyder

## Abstract

Quantification of cellular deoxyribonucleoside mono-(dNMP), di-(dNDP), triphosphates (dNTPs) and related nucleoside metabolites are difficult due to their physiochemical properties and widely varying abundance. Involvement of dNTP metabolism in cellular processes including senescence and pathophysiological processes including cancer and viral infection make dNTP metabolism an important bioanalytical target. We modified a previously developed ion pairing reversed phase chromatography-mass spectrometry method for the simultaneous quantification and ^13^C isotope tracing of dNTP metabolites. dNMPs, dNDPs, and dNTPs were chromatographically resolved to avoid mis-annotation of in-source fragmentation. We used commercially available ^13^C^15^N-stable isotope labeled analogs as internal standards and show that this isotope dilution approach improves analytical figures of merit. At sufficiently high mass resolution achievable on an Orbitrap mass analyzer, stable isotope resolved metabolomics allows simultaneous isotope dilution quantification and ^13^C isotope tracing from major substrates including ^13^C-glucose. As a proof of principle, we quantified dNMP, dNDP and dNTP pools from multiple cell lines. We also identified isotopologue enrichment from glucose corresponding to ribose from the pentose-phosphate pathway in dNTP metabolites.

## Introduction

Intracellular deoxyribonucleoside triphosphate (dNTP) supply is tightly controlled by *de novo* synthesis, salvage, and degradation pathways (1, 2). Aberrant concentrations of dNTPs and their metabolites are associated with control of senescence (3-5), cancer (6, 7), metabolic diseases (3, 8), neurodegeneration (9), and viral infection (10). Basal dNTP levels during G1/G0 are extremely low (femtomolar range) and are mainly used for DNA repair and mitochondrial DNA synthesis (11, 12). During S phase, dNTP levels increase ten-fold to accommodate nuclear DNA replication (13). Excess dNTP levels reduce genome stability, replication fidelity, and reduce the length of, or delay entry into S phase (14). On the other hand, low dNTP levels are deleterious by increasing the erroneous incorporation of rNMPs as well as the incidence of replication stress, leading to fork arrest, collapse, and double strand breaks (15). Mitochondrial depletion also results from the imbalance of dNTP levels (16). Therefore, dNTP pool balance is critical for the health of the cell.

Low basal concentrations of dNTPs and interference from analogs including NTPs, dideoxynucleoside triphosphates and mono‐ or di-phosphate deoxynucleosides presents an analytical challenge (17, 18). Detection with an enzymatic assay, in which a radioactive or fluorescence labeled dNTP is incorporated by a DNA polymerase proportionally to the unknown complementary base, is relatively sensitive with limits of detection (LoD) in the low pmol range (12). DNA polymerases sometimes incorporate rNTPs, thus often overestimating the concentration of dNTPs, especially at low dNTP concentrations or in complex matrices (12, 19). Furthermore, the multiplexing ability of these assays are limited and they cannot simultaneously quantify all nucleosides and their corresponding mono-, di-, and tri-phosphate forms (20). Liquid chromatography (LC)-UV detection methods are useful in dNTP bioanalysis but are limited in sensitivity, specificity, and multiplexing ability (21).

Mass spectrometry (MS) based quantification of dNTP metabolism offers unique benefits in terms of multiplexing, sensitivity and specificity. However, in-source fragmentation in electrospray ionization (ESI) sources of tri-phosphates to mono‐ and di-phosphates is problematic without chromatographic resolution or preparative separation of the mono-, di‐ and tri-phosphates (22). Specifically, this complicates direct infusion high resolution mass spectrometry (HRMS) methods developed for other nucleotide metabolites (23). Triple quadrupole LC-tandem MS (LC-MS/MS) based methods have been developed to provide increased multiplexing and more specific measurements of dNMP and dNTP metabolites at adequate sensitivity for most biological samples (24). Methods for dNDPs are sparse in the literature, despite the biological importance of the generation of these metabolites via ribonucleotide reductase. Alternatively, due to the analytical challenges in quantifying dNTPs, many studies indirectly quantify dNTP metabolism by strategically assaying precursor metabolites including ribose-5-phosphate (R5P) (25).

MS based methods can incorporate isotope dilution by adding an isotope labeled internal standard at the beginning of a bioanalytical workflow to adjust for matrix effects and losses during extraction and analysis (26). ^13^C,^15^N-labeled ribonucleosides are available commercially, but methods to date have not utilized extensive isotope dilution. Aside from use of stable isotopes for more rigorous quantification, MS-based analysis can also enable isotope tracing, where the incorporation of a stable isotope labeled substrate in a metabolically active system is quantified by measuring the incorporation of the isotope label into an analyte. Synthesis of dNTPs incorporate carbon and nitrogen from diverse precursors, in separate pathways for purines and pyrimidines (**Fig. 1**). Carbon sources for *de novo* dNTP synthesis include glucose, aspartate, serine, and glycine whereas nitrogen is derived from glutamine, or aspartate (13). These precursors can be derived from multiple metabolic pathways, including glycolysis, glutaminolysis, and TCA cycle metabolism (3). Preferential utilization of these metabolites through each of these pathways is both cell-type and context dependent. Therefore, tracing the fate of atoms from different metabolites to dNM/D/TP pools is useful for understanding pathophysiological metabolic rewiring and pharmacological intervention.

**Figure 1.**
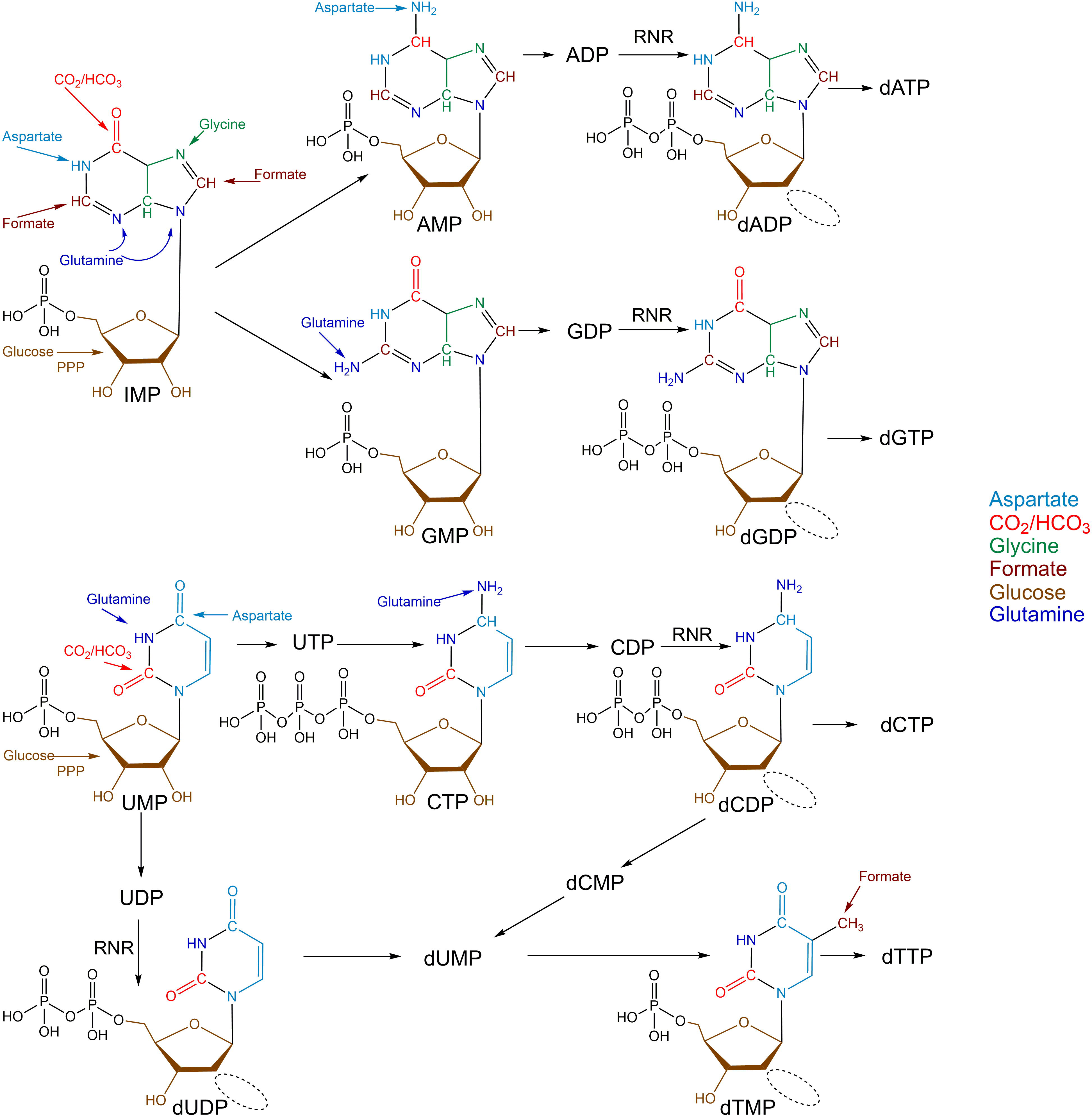
dNTP metabolism integrates atoms from a number of potential substrates.

To overcome the analytical challenges of dNTP quantification and maximize the analytical benefits of MS based analysis, we developed an ion pairing LC-HRMS method for the simultaneous isotope dilution based quantification and ^13^C-isotope tracing of dNTP from major carbon precursors. Mono-, di-, and tri-phosphates of deoxyribonucleosides were quantified across different cell lines, and isotopic incorporation from ^13^C-glucose was examined. We demonstrate improved analytical parameters over label-free quantification and the ability to discern the contribution of major carbon sources into biologically informative patterns of isotope incorporation.

## Materials and Methods

### Chemicals

Standards for dAMP, ADP, dADP, ATP, dATP, dTMP, dTDP, dTTP, GMP, dGMP, GDP, dGDP, GTP, dGTP, CMP, dCMP, CDP, dCDP, CTP, dCTP, UMP, dUMP, UDP were from Sigma-Aldrich (St Louis, MO). Stable isotope labeled internal standards AMP-^13^C_10_,^15^N_5_, dAMP-^13^C_10_,^15^N_5_, ATP-^13^C_10_,^15^N_5_, dATP-^13^C_10_,^15^N_5_, dTMP-^13^C_10_,^15^N_2_, dTTP-^13^C_10_,^15^N_2_, GMP-^13^C_10_,^15^N_5_, dGMP-^13^C_10_,^15^N_5_, GTP-^13^C_10_,^15^N_5_, dGTP-^13^C_10_,^15^N_5_, dCMP-^13^C_9_,^15^N_3_, CTP-^13^C_9_,^15^N_3_, dCTP-^13^C_9_,^15^N_3_, UTP-^13^C_9_,^15^N_2_, CMP-^13^C_9_,^15^N_3_, UMP-^13^C_9_,^15^N^2^ were also from Sigma-Aldrich. No suitable source of dUDP or stable isotope labeled ADP, dADP, dTDP, GDP, dGDP, CDP, dCDP, dUMP, UDP, dUDP, or dUTP was found.

Diisopropylethylamine (DIPEA) and 1,1,1,3,3,3-hexafluoro 2-propanol (HFIP), were purchased from Sigma-Aldrich. Optima LC-MS grade water, methanol, and acetonitrile (ACN) were purchased from Thermo Fisher Scientific (Waltham, MA).

### Cell culture

Normal, diploid IMR90 human fibroblasts were cultured according to the ATCC in low oxygen (2%) in DMEM (4.5 g/L glucose, Corning cat#10017CV) with 10% FBS supplemented with L-glutamine, non-essential amino acids, sodium pyruvate, and sodium bicarbonate. Experiments were performed on IMR90 between population doubling #25-35. Cells were routinely tested for mycoplasma as described in (27). Experiments targeting Ribonucleotide Reductase Regulatory Subunit M2 (RRM2), p16, mammalian target of rapamycin (mTOR), and ATR serine/threonine kinase (ATR) were previously included in Buj, *et al*.(28). Lentiviral constructs were transfected into 293FT cells using Lipofectamine 2000 (Thermo Fisher, Cat# 11668019). Lentivirus was packaged using the ViraPower Kit (Invitrogen, cat# K497500) following the manufacturer’s instructions. Briefly, IMR90 were infected with pLKO.1 or pLKO.1-shRRM2 (Sigma-Aldrich, TRCN0000049410) and 24 hours later cells were infected with pLKO.1 or pLKO.1-shp16 (Sigma-Aldrich, TRCN0000010482). Cell were selected with puromycin (3 µg/mL) for seven days. Cells were treated at day 4 with 0.5nM of Temsirolimus an mTOR inhibitor (Selleckchem cat#S1044) or VE822 an ATR inhibitor (cat#S7102). After that, media was replaced for DMEM without glucose and glutamine (Sigma-Aldrich, cat#D5030) and supplemented with 0.5% of charcoal stripped FBS (Sigma-Aldrich, cat#F6765-500ML), 20mM HEPES (Fisher BioReagents, cat# BP310) and 5mM of [^13^C_6_]-D-glucose (Sigma-Aldrich, Cat# 389374) or natural isotopic abundance D-glucose (Sigma-Aldrich, cat#G8644). Labeling was stopped after 8 hours. Cells were washed twice with PBS and harvested with trypsin (Corning cat#25-053-Cl). Pelleted cells were snap frozen in liquid nitrogen and stored at −80 Celsius. Cell pellets were extracted before analysis by addition of a 50 µL (20 ng/µL) mix of all stable isotope labeled internal standards (1000 ng/sample) in 80:20 (v/v) methanol:water followed by 1 mL of −80°C 80:20 (v/v) methanol:water.

### Liquid chromatography-high resolution mass spectrometry

LC-HRMS was as previously described with minor modifications (29). Briefly, an Ultimate 3000 UHPLC equipped with a refrigerated autosampler (at 6 °C) and a column heater (at 55 °C) with a HSS C18 column (2.1 × 100 mm i.d., 3.5 μm; Waters, Milford, MA) was used for separations. Solvent A was 5 mM DIPEA and 200 mM HFIP and solvent B was methanol with 5 mM DIPEA and 200 mM HFIP. The gradient was as follows: 100 % A for 3 min at 0.18 mL/min, 100 % A at 6 min with 0.2 mL/min, 98 % A at 8 min with 0.2 mL/min, 86 % A at 12 min with 0.2 mL/min, 40 % A at 16 min and 1 % A at 17.9 min-18.5 min with 0.3 mL/min then increased to 0.4 mL/min until 20 min. Flow was ramped down to 0.18 mL/min back to 100 % A over a 5 min re-equilibration. For MS analysis, the UHPLC was coupled to a Q Exactive HF mass spectrometer (Thermo Scientific, San Jose, CA, USA) equipped with a HESI II source operating in negative mode. The operating conditions were as follows: spray voltage 4000 V; vaporizer temperature 200 °C; capillary temperature 350 °C; S-lens 60; in-source CID 1.0 eV, resolution 60,000. The sheath gas (nitrogen) and auxiliary gas (nitrogen) pressures were 45 and 10 (arbitrary units), respectively. Single ion monitoring (SIM) windows were acquired around the [M-H]^−^ of each analyte with a 20 *m/z* isolation window, 4 *m/z* isolation window offset, 1e^6^ ACG target and 80 ms IT, alternating with a Full MS scan from 70-950 *m/z* with 1e6 ACG, and 100 ms IT.

### Statistical analysis

Data was analyzed in XCalibur v4.0 and/or Tracefinder v4.1 (Thermo). Statistical analysis and graph generation was conducted in GraphPad Prism v7.04 (GraphPad Software La Jolla, CA).

## Results/Discussion

### Ion pairing reversed phase separation allows chromatographic resolution of intact mono-, di‐ and tri-phosphate forms of nucleosides

Tri‐ and di-phosphates are known to easily lose a phosphate group in the ESI source to generate di‐ and mono-phosphates (22). Therefore, it is critical that the mono-, di-, and tri-phosphates of dNTP metabolites are resolved by chromatography. We first determined retention times and confirmed high abundance ions by injection of each individual standard. Mono-phosphates eluted in the 4-8 minute range, di-phosphates in 12-14.5 and triphosphates from 15-onwards. This provided baseline resolution of the major in-source fragments that had the same retention time of the precursor tri‐ or diphosphate. Standard for dAMP was cross-contaminated with other NTP metabolites and were not included in our analysis for purposes of absolute quantification. To confirm that this method was adequate for detection in cell samples, methanolic extract of IMR90 cells was analyzed and extracted ion chromatograms for all analytes and their available stable isotope labeled internal standards were plotted (**Fig. S1**). For analytes without matched isotope labeled analogs, the identity of peaks were confirmed by co-elution with pure standards to ensure mis-identification with the in source fragment of related analytes (**Fig. S2**). Resolution of the in-source fragments was maintained in cell matrix. Operation of the HRMS in full scan alone was insufficiently sensitive to detect all dNTP metabolites, thus in the final method single ion monitoring (SIM) was used, with a 20 *m/z* window with an offset of 4 *m/z* around each *m/z* corresponding to each analyte [M-H]^−^. This window includes all internal standards used, and allowed for simultaneous mass isotopologue distribution analysis described below.

We were not able to chromatographically baseline resolve the isobaric dGTP/ATP, dGDP/ADP, dGMP/AMP pairs in our system. Overlap of these nucleotides was noted in other methods, and negative mode fragmentation was reported to be less specific than positive ion mode. In the cases of both dGM/D/TPs, we could resolve the deoxynucleotides with LC-MS/HRMS, but at drastically reduced sensitivity with more limited and complicated isotopologue analysis since the entire nucleotide is not captured by the detected ions (data not shown). Thus, we excluded this from analysis in this method. The concentration of AMP/ADP/ATP is expected to be orders of magnitude higher than the deoxyguanosine metabolites; thus, quantification by this method would reflect by majority AMP/ADP/ATP.

### Stable isotope dilution improves analytical performance of LC-HRMS based quantification of dNTPs

Standard curves were generated for dNTPs except dAMP and dGM/D/TP. Pilot testing indicated that using higher amounts of dNTP internal standards improved detection in the lower range of standard curves, without interference from residual unlabeled impurities. Analysis of curves for dTM/D/P was conducted with isotope dilution or by label-free quantification using peak area around the levels detected in cell samples. For isotope dilution, the simplest model that fit the data with linear least-squares regression around the range detected in 1×10^6^ cells (0.97-500 ng/sample) was used, with R^2^ values of 0.9979 (no weighting), 0.9973 (1/x weighting), and 0.9933 (1/x weighting) for dTMP, dTDP, and dTTP, respectively. In comparison with the same weightings, label-free quantification gave R^2^ values of 0.9977, 0.8258, and 0.8592 (**Fig. 2 A,B**).

**Figure 2.**
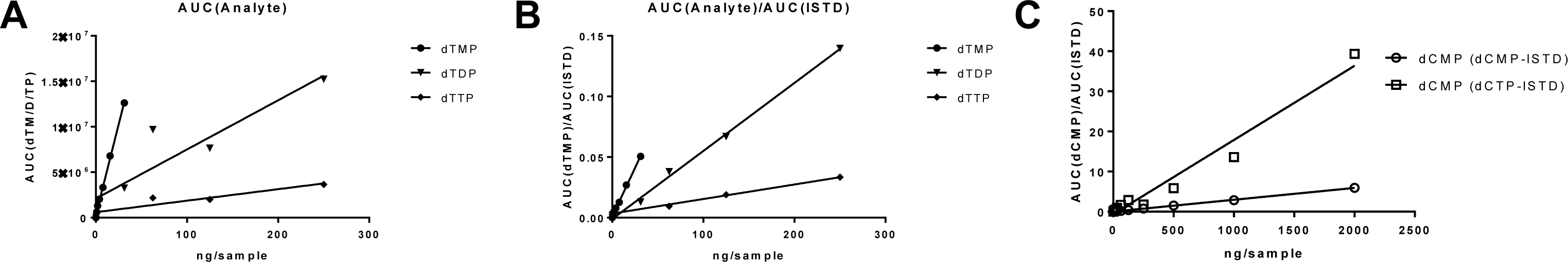
Calibration curves of dTM/D/P in cell samples using A) label-free quantification B) isotope dilution. C) Calibration curves of dCMP using dCMP-internal standard (ISTD) or dCTP-ISTD.

Calibration curves for dCM/D/TP were also linear across the range encountered in cell samples with excellent R^2^ values of 0.9999, 0.9968, 0.9965 for dCMP, dCDP and dCTP respectively. Since previous methods have used surrogate internal standards (most commonly using 1 or two internal standards to normalize multiple analytes), we tested the utility of dCMP analysis by normalization to either the matched dCMP analog or dCTP. Normalization of dCMP signal area under the curve by dCTP-internal standard area under the curve yielded a poor least squares fit at the same concentration ranges (**Fig. 2 C**).

The signal intensity in the blank containing only isotope labeled internal standards had consistent background with 0 intensity. Therefore, for purposes of comparison to other methods of the limit of quantitation (LoQ), we conservatively estimated the LoQ as the first point that would fall within the experimentally confirmed linear range of the method at around 0.03 pmol (30 fmol) on column for dTMP, dTDP and dTTP. This was more than sufficient to quantify the levels within cells lines tested.

Quality controls were created to bracket levels observed in cell studies and from remnant tumor and non-tumor pathologic tissue (**Table 1**). This ranged from 1 ng/sample to 600 ng/sample with pmol amounts reported for on column and pmol per sample for the lowest quality control.

**Table 1.**
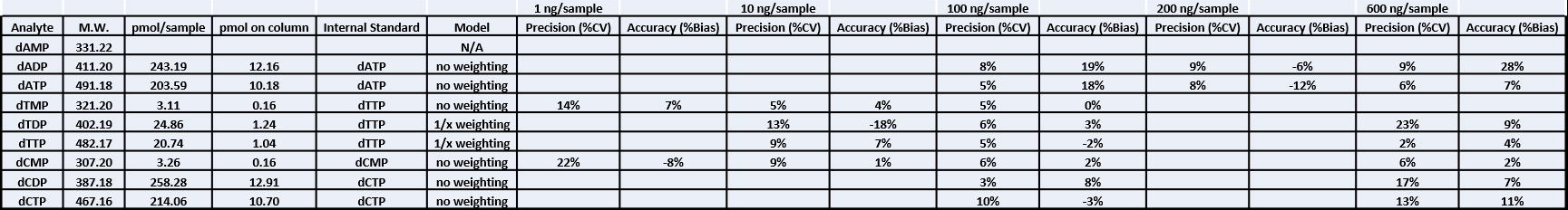
Analytic figures of merit for quality control samples bracketing cell and tissue levels of dNM/D/TPs.

|  |  |  |  |  |  | 1 ng/sample |  | 10 ng/sample |  | 100 ng/sample |  | 200 ng/sample |  | 600 ng/sample |  |
| --- | --- | --- | --- | --- | --- | --- | --- | --- | --- | --- | --- | --- | --- | --- | --- |
| Analyte | M.W. | pmol/sample | pmol on column | Internal Standard | Model | Precision (%CV) | Accuracy (%Bias) | Precision (%CV) | Accuracy (%Bias) | Precision (%CV) | Accuracy (%Bias) | Precision (%CV) | Accuracy (%Bias) | Precision (%CV) | Accuracy (%Bias) |
| dAMP | 331.22 |  |  |  | N/A |  |  |  |  |  |  |  |  |  |  |
| dADP | 411.20 | 243.19 | 12.16 | dATP | no weighting |  |  |  |  | 8% | 19% | 9% | -6% | 9% | 28% |
| dATP | 491.18 | 203.59 | 10.18 | dATP | no weighting |  |  |  |  | 5% | 18% | 8% | -12% | 6% | 7% |
| dTMP | 321.20 | 3.11 | 0.16 | dTTP | no weighting | 14% | 7% | 5% | 4% | 5% | 0% |  |  |  |  |
| dTDP | 402.19 | 24.86 | 1.24 | dTTP | 1/x weighting |  |  | 13% | -18% | 6% | 3% |  |  | 23% | 9% |
| dTTP | 482.17 | 20.74 | 1.04 | dTTP | 1/x weighting |  |  | 9% | 7% | 5% | -2% |  |  | 2% | 4% |
| dCMP | 307.20 | 3.26 | 0.16 | dCMP | no weighting | 22% | -8% | 9% | 1% | 6% | 2% |  |  | 6% | 2% |
| dCDP | 387.18 | 258.28 | 12.91 | dCTP | no weighting |  |  |  |  | 3% | 8% |  |  | 17% | 7% |
| dCTP | 467.16 | 214.06 | 10.70 | dCTP | no weighting |  |  |  |  | 10% | -3% |  |  | 13% | 11% |

To examine the effect of isotope dilution versus label free quantification we analyzed data from IMR90 cells with various short hairpin knockdown of genes found to disrupt nucleotide metabolism (28). This provided a bioanalytical meaningful way to modulate dNTP metabolites within similar cellular backgrounds. For this analysis, the analyst was blinded to identity of each sample (with respect to control versus knockdown and inhibition). Comparison of values obtained from quantification from cells revealed better precision with isotope dilution for dTMP (**Fig. 3 A,D**), dTDP (**Fig. 3 B,E**), and dTTP (**Fig. 3 C,F**). Bland-Altman plots showing the % difference between the estimated value by each method and the averages of both approaches revealed an interesting concentration dependent bias in in all three analytes, with the lower values exhibiting a stronger negative bias in label-free quantification. This bias was especially pronounced in the IMR90 cell lines which had the three lowest values of all dTM/D/TP metabolites. Label-free quantification under-estimated the pools in IMR90 by around 175%, 90%, and 50% for dTMP, dTDP and dTTP. The reasons for this differential bias across nucleotide amounts is unknown, but has implications for the reporting of nucleotide metabolites from metabolomics data.

**Figure 3.**
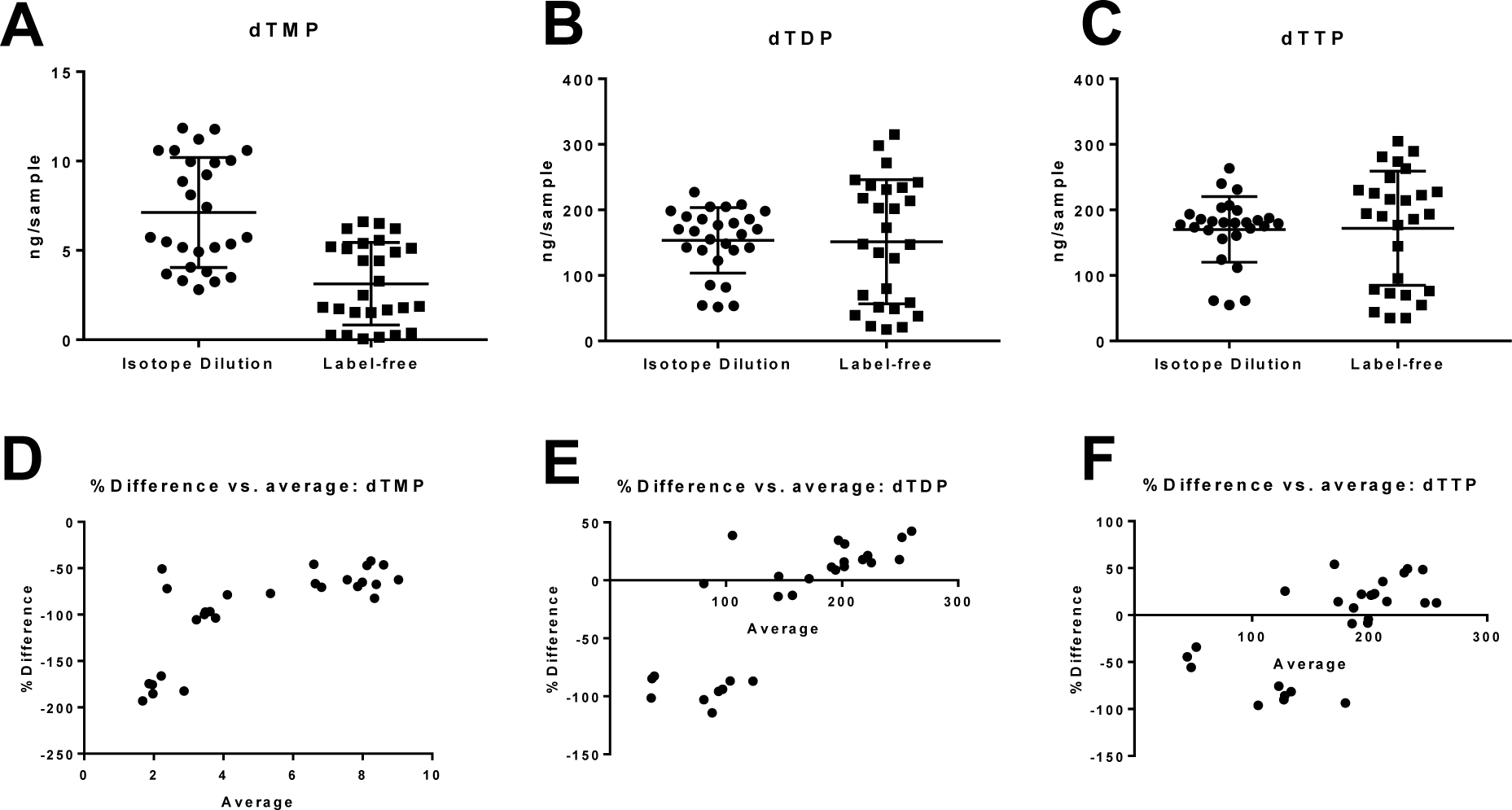
Bias in quantification of dTM/D/TP in cell samples using isotope dilution or label-free quantification. Quantification of A) dTMP B) dTDP C) dTTP. Bland-Altman plot showing % difference in isotope dilution versus label-free quantification for D) dTMP E) dTDP and F) dTTP.

### Stable isotope resolved metabolomics of dNTPs is possible at ultra-high resolution

Carbon atoms are derived from glucose, CO_2_/bicarbonate, formate, aspartate, and glycine, with nitrogen atoms coming from glutamine and aspartate. Since none of these are essential amino acids or nutrients they can be derived from the diverse sources of all of these precursors as well. These precursors can be traced via stable isotope labeling strategies which incorporate atoms from a labeled substrate into the metabolic product. Isotopic tracing requires higher sensitivity than quantification as isotopologue analysis requires, by definition, analysis of less abundant isotopologues. Furthermore, tracing with a stable isotope splits the signal intensity across isotopologues as they incorporate the isotopic label. Theoretically, at sufficient mass resolution, simultaneous quantification and isotope tracing can be accomplished with differential neutron encoded labels (e.g., ^13^C vs. ^15^N) due to the mass defect in the additional neutron in atoms with varying nuclei (30).

At sufficient mass resolution, stable isotopes of commonly used tracing atoms (_2_H, ^13^C, ^15^N, ^18^O, etc.) can be resolved from each other. Thereby, neutron-encoded information enables simultaneous metabolic tracing by stable isotope labeling with orthogonal stable isotope labeling used as an internal standard for isotope dilution based quantification (30). Since dNTPs and nucleoside metabolites incorporate C, H, N, and O atoms, and commercially available ^13^C,^15^N-labeled pure standards are available for isotope dilution, we examined the ability to simultaneously trace ^13^C-labeling via metabolic tracing with ^13^C_6_-glucose with ^13^C, ^15^N internal standards. This allows an experimental design where the origin of dNTP pools can be quantitatively examined, by quantifying the substrate-product relationship from labeled precursors, as well as the relative importance of *d*e *novo* versus different salvage or uptake pathways. The limiting factor is resolution of the ^13^C and ^15^N labels, so we modeled theoretical resolution at different resolving powers. With Orbitrap mass analyzers, resolution falls off with increasing *m/z* (31), thus the most difficult to resolve isotopologues are the triphosphates with the highest possible number of labels. 240,000 resolving power was sufficient to baseline resolve the ^15^N_1_ labeled isotopologue from ^13^C_1_ labels (**Fig. 4A-D**). A lower resolution setting would be capable of resolving the dNMPs, or utilization of MS/HRMS and analysis of the dNMP product ions (**Fig. 4E**).

**Figure 4.**
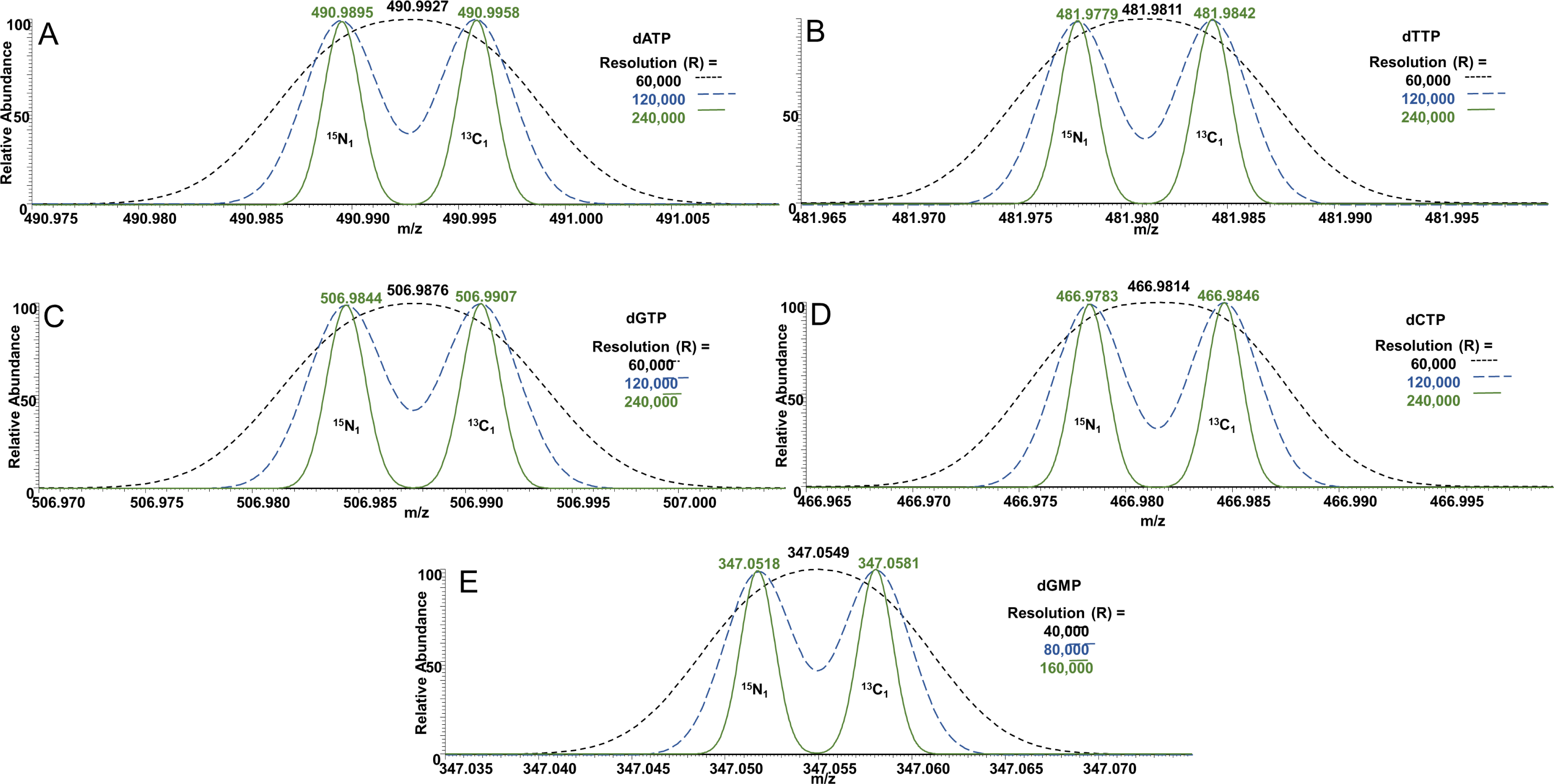
Resolution of the dNTPs and their ^13^C/^15^N isotopologues requires ultra-high resolution mass spectrometry. Theoretical Gaussian resolution (FWHM) with 40 samples per peak is shown at increasing levels of resolving power per nucleotide. Predicted centroid masses for [M-H]^−^at each resolution are shown for a 1:1 mixture each of (A) ^13^C_1_dATP/^15^N_1_dATP (B)^13^C_1_dTTP/^15^N_1_dTTP (C) ^13^C_1_dGTP/^15^N_1_dGTP (D) ^13^C_1_dCTP/^15^N_1_dCTP and (E) ^13^C_1_dGMP/^15^N_1_dGMP.

We tested this theoretical possibility by a proof-of-principle experiment using isotope dilution absolute quantification in the analysis of a ^13^C_6_-glucose labeling experiment. IMR 90 cells were grown with or without ^13^C_6_-glucose, and with either empty vector control or RRM2 knockdown. We analyzed the dTM/D/TP pool size by isotope dilution based quantification (**Fig. 5 A**) and the isotopologue enrichment of each analyte (**Fig. 5 B,C,D**). We integrated the peak area of each isotopologue of dTM/D/TP corresponding to the sequential incorporation of ^13^C atoms. After correction for natural isotopic abundance isotopologue enrichment of dTM/D/TP revealed the quantitative incorporation of ^13^C labeled glucose in a pattern of mixed sequential M1 and M2 labeling with a large increase in M5 labeling and some M6 labeling. This peak at M5 likely corresponds to the incorporation of ^13^C_5_.

**Figure 5.**
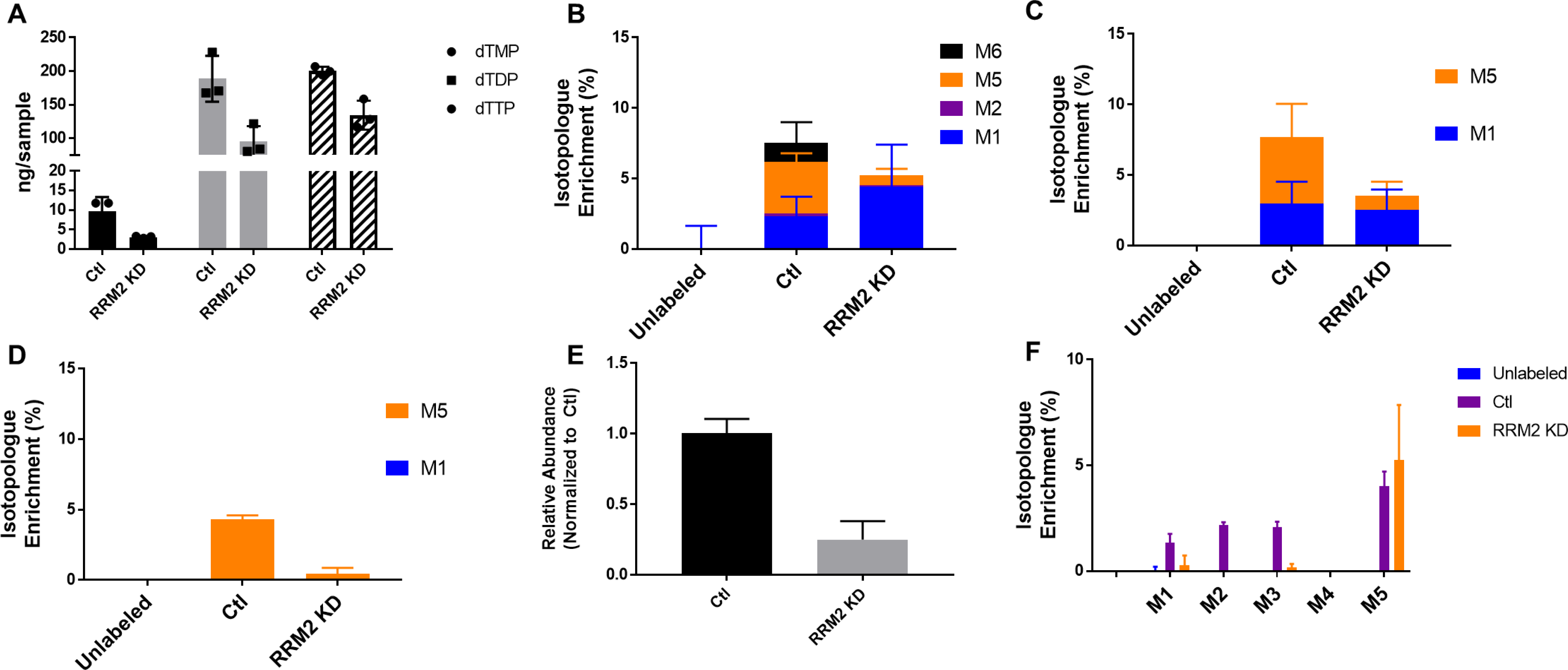
Simultaneous isotope dilution based quantification and isotope tracing of dTM/D/P in ^13^C labeled IMR90 extract. A) Quantification of dTM/D/TP, isotopologue enrichment (%) of B) dTMP C) dTDP and D) dTTP. Full scan data was integrated for E) relative quantification of ribose-5-phosphate (R5P) F) isotopologue enrichment (%) of R5P.

Since full scan data was acquired in addition to the targeted SIMs, we re-interrogated the data for relative abundance and labeling into ribose-5-phosphate, the pentose-phosphate pathway product that provides the ribose in dNTPs (**Fig. 5 E,F**). Labeling of ribose-5-phosphate mirrored that of dNTPs, providing confirmation that we were measuring the incorporation of the ^13^C into dNTPs. Recent work has demonstrated the importance of nucleotide tracing. For instance, that response of cancer cells to metformin correlates with nucleotide levels derived from TCA carbon sources but not the pentose phosphate pathway (32). Similarly, resistance to the antimetabolite gemcitabine correlates with glucose carbon flux specifically through the non-oxidative pentose phosphate pathway (33). Thus, tracing of metabolites to nucleotides/dNTPs may determine which patients will respond to metabolic inhibitors and reveal metabolic vulnerabilities for subsets of cancers. However, many studies have not confirmed tracing of metabolites from nucleotide precursors to dNTPs. It will be important in the future to further dissect the regulation of dNTP synthesis using tracing studies in both normal and diseased states.

There are two major caveats of this method. First, further improvement of this method could be made by chromatographically resolving the remaining isobars (AMP/dGMP, ADP/dGDP, ATP/dGTP). Although we could find a specific fragment on tandem MS/HRMS to differentiate the adenosine nucleotide phosphates from the deoxyguanosine nucleotide phosphates, this reduced sensitivity of the method below that useful for our purposes. Second, although we did not quantify it due to an impurity in the dAMP standard, sensitivity of dAMP was reduced by a high-intensity contaminant ion present across the LC-gradient falling in the same SIM window. This is problematic on our Orbitrap instrument as the sensitivity benefit gained from using the quadrupole for isolation to restrict ions allowed into the C-trap and Orbitrap is negated by a high-intensity background ion within the SIM window when the quadrupole isolation window was sufficient to allow complete isotopologue enrichment analysis.

## Conclusion

Direct quantification and isotope tracing of dNTP metabolites is analytically challenging. Ion pairing reversed phase LC coupled with Orbitrap based high-resolution mass spectrometry provides a platform for simultaneous isotope dilution based quantification of dNTPs with isotope tracing from multiple potential substrates.

## Acknowledgements

This work was made possible by a NARSAD young investigator grant from the Brain and Behavior Foundation (NWS) as well as NIH grants K22ES026235 (NWS), R00CA194309 (KA), and P30ES013508.

---

**Supplemental Figure 1.**
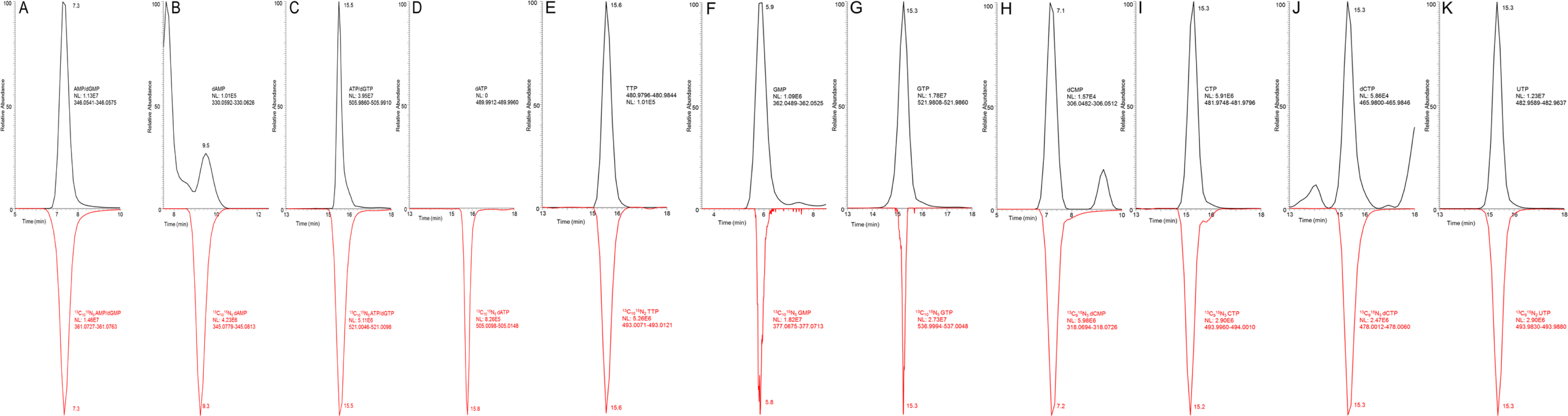
Extracted ion chromatograms of deoxynucleotides with their stable isotope labeled analogs.

**Supplemental Figure 2.**
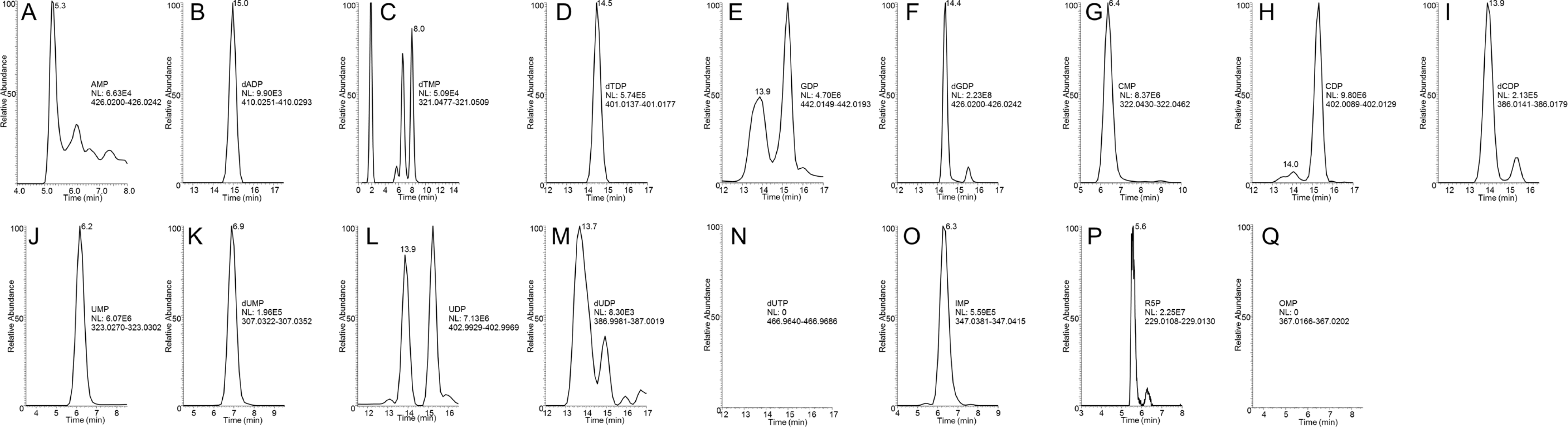
Extracted ion chromatograms of deoxynucleotides. Where multiple peaks are observed in the chromatogram, the correct peak as determined by co-elution with a pure standard are indicated by labeling the retention time of the empirically assigned peak.

